# Population-based metaproteomics reveals functional associations between gut microbiota and phenotypes

**DOI:** 10.1101/2024.11.26.625569

**Authors:** Shuang Liang, Yingying Sun, Zelei Miao, Bang-yan Li, Ziyuan Xing, Yuting Xie, Enci Cai, Sainan Li, Pu Liu, Min Yang, Menglei Shuai, Wanglong Gou, Wenhao Jiang, Youming Wang, Huanhuan Gao, Ke Zhang, Jing Yu, Xue Cai, Xingbing Wang, Yu-ming Chen, Ju-Sheng Zheng, Tiannan Guo

**Affiliations:** Affiliated Hangzhou First People’s Hospital, State Key Laboratory of Medical Proteomics, School of Medicine, Westlake University, Hangzhou, Zhejiang Province, China; Westlake Center for Intelligent Proteomics, Westlake Laboratory of Life Sciences and Biomedicine, Hangzhou, Zhejiang Province, China; Research Center for Industries of the Future, School of Life Sciences, Westlake University, Hangzhou, Zhejiang Province, China; State Key Laboratory for Managing Biotic and Chemical Treats to the Quality and Safety of Agro-products, Zhejiang Academy of Agricultural Sciences, Hangzhou 310021, China; Zhejiang Key Laboratory of Multi-Omics in Infection and Immunity, Center for Infectious Disease Research, School of Medicine, Westlake University, Hangzhou, China; Department of Epidemiology, Guangdong Provincial Key Laboratory of Food, Nutrition and Health, School of Public Health, Sun Yat-sen University, Guangzhou, China; Westlake Omics (Hangzhou) Biotechnology Co., Ltd., No. 1 Yunmeng Road, Cloud Town, Xihu District, Hangzhou 310024, Zhejiang Province, China; First Affiliated Hospital of USTC, Division of Life Sciences and Medicine, University of Science and Technology of China, Hefei, Anhui 230001, China

**Keywords:** Metaproteomics, population-based cohort study, human gut microbiome, diabetes, *Megasphaera elsdenii*, butyrate

## Abstract

The association between gut microbiota and physiological disorders underscores the crucial need to understand its largely unexplored biological functions. In this study, we analyzed 86,876 microbial proteins and 1,277 human proteins from 1,967 Chinese stool samples. Our results indicate that metaproteomic data captures the microbial biodiversity of a population more effectively than metagenomic data, displaying greater variability in microbial functions and providing reliable assessments of microbiota stability. We identified crucial functions of core microbes and discovered 10,714 associations between metaproteomic taxon, microbial function, or human proteins with 39 phenotypes, including 1,381 features related to Type 2 Diabetes (T2D). Importantly, *Megasphaera elsdenii*’s conversion of lactate to butyric acid appears to lower blood glucose levels, suggesting a protective mechanism against T2D. Our study underscores the power of population-based metaproteomics to offer unique functional insights into the gut microbiota.

**Highlights:**

- The first large-scale population-based metaproteomics landscape.
- Metaproteomics efficiently maps microbial functions, exposing individual functional variation.
- 10,714 associations between metaproteomic taxa, microbial functions, or human proteins with 39 phenotypes.
- *Megasphaera elsdenii* protects against elevated blood glucose levels in the host by degrading lactate to produce butyric acid.

## Introduction

Apart from their well-known roles in food digestion and nutrition absorption, gut microbes are also involved in multiple physiological processes and disorders including metabolic disorders and cancers^1,2^. The gut microbiota serves as intricate communities, and multi-omic technologies are extensively employed to gauge the dynamic fluctuations within these communities. Studies investigating the relationship between gut microbes and factors such as age, sex, diet, geography, heredity, and health status in specific populations have primarily relied on microbial composition analysis using 16S sequencing or metagenomics^3–7^. Metaproteomics has emerged as a powerful tool to explore the functional diversity of the microbiome, as proteins directly govern cellular functions^8,9^. It can reveal taxonomic biomass contribution variations by quantifying taxon-specific proteins^10^. Additionally, metaproteomics also provides information on human enterocyte proteins, facilitating the investigation of host-microbiome interactions^11,12^. Metaproteomics have been successfully employed in studies of inflammatory bowel disease, diabetes, obesity, and colorectal cancer^13–20^. However, these studies have been restricted to dozens to hundreds of samples primarily due to the relatively long MS time required by the data-dependent acquisition mass spectrometry (DDA-MS) and sophisticated data analysis^21–27^. Therefore, metaproteomics faces challenges in scalability for population-based metaproteome analysis. Due to the highly heterogeneous composition of gut microbiota across individuals^28^, more subjects need to be included to obtain confident biological conclusions. Moreover, no study has systematically investigated the unique insights provided by metaproteomics compared to metagenomics.

Data-independent acquisition (DIA) MS coupled with targeted data analysis enables high-throughput profiling of proteomes^29–32^. In particular, DIA leverages DDA to construct a spectral library, facilitating the precise quantification of thousands of proteins with a single injection. Integrating ion mobility with MS provides additional information into ion size, shape, and charge distribution, thereby enhancing ion separation and reducing chemical noise in measurements^33,34^. As a refinement of DIA, parallel accumulation–serial fragmentation combined with data-independent acquisition (diaPASEF) incorporates trapped ion mobility into the ion separation, leading to a significant improvement in proteomic depth^35,36^. In this study, we performed a diaPASEF-based metaproteomic analysis on a cohort comprising thousands of individuals. The distribution of age, gender, lifestyle, and metabolic diseases in this cohort was representative of the middle-aged and elderly Chinese population. Our findings from population-scale gut metaproteomics offer unique functional insights.

## Results

### Metaproteomic profiling of GNHS cohort

Our study included 1,399 middle-aged and elderly participants from the Guangzhou Nutrition and Health Study (GNHS). Fecal specimens were collected from these individuals, with 568 participants providing samples at baseline and follow-up time points, separated by a median interval of 3.2 years, over a period ranging from 1.4 to 5.5 years. The remaining 831 participants provided samples only once. Altogether, we collected 1,967 fecal samples, each with comprehensive metadata recorded at the time of sampling (Fig. 1A). The metadata comprised 49 clinical and questionnaire-based phenotypes related to lifestyle, demographics, diet, biomedical measures, anthropometrics, metabolic disorders, self-reported diseases, medicine usage, and antibiotic usage (Fig. 1A). Details of these phenotypes are listed in Table S1.

**Figure 1.**
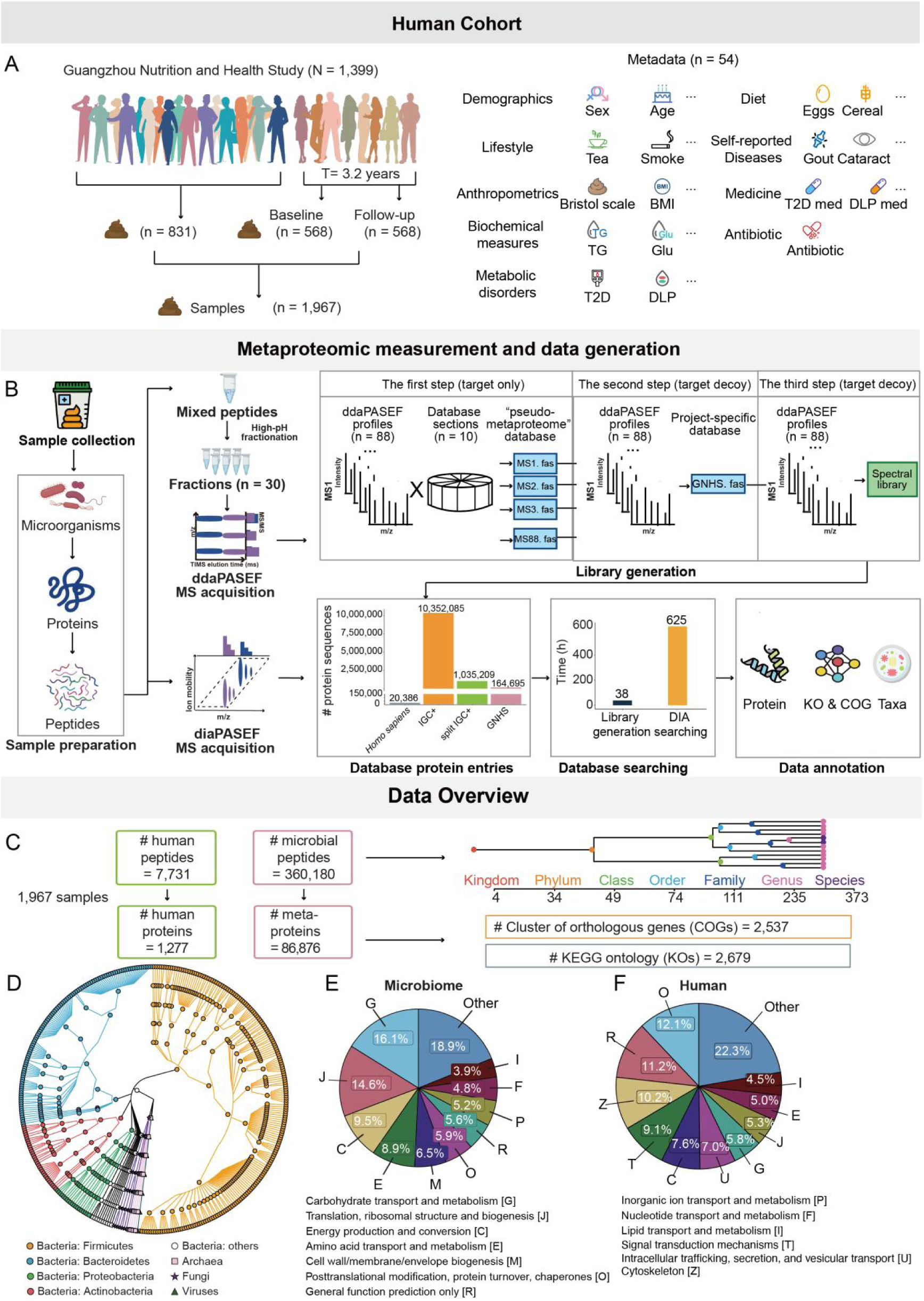
Metaproteomic profiling of the gut microbiome. (A) Population distribution of individuals. Our study covered 1399 individuals from the GNHS cohort. Of these, 831 individuals contributed one sample only, while 568 individuals provided two samples taken approximately 3.2 years apart. This cohort also had 49 available phenotype-related information, including types of sociodemographics, lifestyle, metabolic disorders, biochemical measures, and anthropometrics. (B) The experimental strategy of diaPASEF-based metaproteomic profiling of 2,514 MS runs in the GNHS cohort, of which 1,967 were for our investigations and 547 for quality assessment. (C) Metaproteomic identifications of 1967 fecal samples at the levels of microbial protein, taxonomy, and function. (D) Taxonomic tree based on metaproteomic data. The taxonomic levels range from superkingdoms to species from the inner to the outer circle. (E) The functional distributions of microbiota. The functional category was annotated using the COG database. (F) The functional distributions of human proteins. The functional category was annotated using the COG database.

We employed the diaPASEF-based metaproteomic analysis to study the functional profiles of gut microbiota in a cohort of thousands of individuals. A total of 2,514 MS profiles containing 1,967 fecal samples and 547 quality controls were acquired. Our investigation encountered an initial hurdle in the analysis of mass spectral data. This challenge is commonplace in metaproteomic analysis, primarily due to the extensive protein sequence databases inherent to the human gut microbiome, necessitating significant computational resources. Additionally, the integration of ion mobility dimensions in diaPASEF technology introduces further complexity to the data, rendering its interpretation more intricate. To surmount these obstacles, we developed a customized analysis pipeline specifically designed for diaPASEF-based metaproteomics. This pipeline encompasses ddaPASEF-based spectral library generation, database searches tailored to diaPASEF MS profiles, and the annotation of taxonomic and functional information within metaproteomic datasets. For the spectral library generation, we employed a refined three-step iterative database search method^37^ alongside protein inference rules based on the maximum parsimony^38^ (Fig. 1B, Fig. S1, Fig. S2). The process commenced with partitioning the extensive human gut protein sequence database into ten segments, each dedicated to the target-only proteomic search of a ddaPASEF MS profile. The identified protein sequences obtained from the target-only search were merged to create a “pseudo-metaproteome” database for each ddaPASEF MS profile (Fig. 1B, Fig. S1). Subsequently, a project-specific database (GNHS.fas) was constructed by matching each profile against its corresponding “pseudo-metaproteome,” utilizing a target-decoy approach. This database, comprising proteins identified with high confidence (FDR ≤ 0.01) from all ddaPASEF analyses, served to pinpoint project-specific microbial proteins accurately. The final phase involved searching all ddaPASEF profiles against this project-specific database through standard target-decoy search strategies. Subsequently, an in-house protein inference process based on the “maximum parsimony principle” was applied to eliminate the redundant proteins (Figure S2 and methods), resulting in the construction of the definitive spectral library.

Employing a three-step iterative search strategy enabled us to notably diminish the size of the protein sequence database. Initially, database segmentation reduced the entry count from 10,352,085 to 1,035,209. Subsequent refinement, incorporating metaproteomic data from GNHS population, further reduced this to just 164,695 entries, denoted as “GNHS.fas”. This streamlined process facilitated the construction the spectral library within a timeframe of 38 hours. The subsequent protein characterization and quantification via diaPASEF MS-based analysis of all samples required approximately 625 hours, averaging about 15 minutes per MS profile (Fig. 1B). Finally, taxonomic and functional annotations were performed leveraging the clusters of orthologous genes (COG) and KEGG ontology (KO) databases, augmenting our metaproteomic data with significant biological insights.

### Data overview

For all the 1,967 fecal samples, we measured 7,731 human-specific peptides belonging to 1,277 unique human proteins, 360,180 microbial-specific peptides belonging to 86,876 microbial proteins (Fig. 1C). Of all the microbial proteins, 74,624 are unique microbial proteins and 12,252 are microbial protein groups comprising undistinguishable microbial proteins. Here, we refer to the unique microbial proteins and the microbial protein groups as “metaproteins”. For the taxonomic characterization, we identified four superkingdoms, 34 phyla, 49 classes, 74 orders, 111 families, 235 genera, 373 species, and 146 strains. For the functional characterization, a total of 2,537 COGs and 2,679 KO clusters were annotated (Fig. 1C). To assess the stability of human protein detection, we subjected the fecal samples to multiple washes, ranging from three to five times. Remarkably, a significant majority (86.3%) of these human proteins were consistently detected after multiple washes, underscoring their robustness and stability in the human fecal samples (Fig. S3 A-E). Further Gene Ontology (GO) analysis revealed an enrichment of these proteins in peptidase activity, highlighting their pivotal role in food digestion processes (Fig. S3F). Additionally, 96 human proteins have been shown to interact with gut microbes and are potentially implicated in regulating human diseases ^39^ (Fig. S3G). Furthermore, to assess data quality over extended acquisition periods, we obtained multiple pairs of technical and biological replicate samples within and among batches. We observed high correlation and closely clustered Bray-Curtis (BC) distance among the 547 QC replicates, which included 95 inter-batch biological replicates, 98 inter-batch technical replicates, 169 intra-batch biological replicates, and 185 intra-batch technical replicates (Fig. S4A). The BC distance across these replicates showed no significant batch-to-batch variances (Fig. S4B). Principal Component Analysis (PCA) further validated our results, demonstrating a consistent distribution of samples with minimal batch effects, thus affirming the integrity and reliability of the data (Fig. S4C).

In each sample, we identified an average of 477 ± 114 human proteins (mean ± SD), 9,761 ± 3,905 metaproteins, 103 ± 18 genera, 140 ± 29 species, 1,304 ± 268 KO, and 1,365 ± 241 COGs clusters. The median taxonomic annotation rate per sample was 86.0%, while the median functional annotation rates per sample were 75.2% for KO and 93.9% for COG clusters (Fig. S5 A–D). A considerable metaproteome variation in richness was observed on the microbial function and taxonomy levels (Fig. S5E), consistent with findings from a previous population-scale metagenomic study^4^. Based on the metaproteomic data, we gained a fundamental understanding of the taxonomy and functions of the gut microbiome across the GNHS population (Fig. 1 D–E, Fig. S6). The dominant bacterial phyla were Firmicutes, Bacteroidetes, Proteobacteria, and Actinobacteria (Fig. 1D, Fig. S6A), which aligns with the finding of a recent population-scale metagenomic study ^40^. Apart from Bacteria, we also identified six microbes from Archaea and Eukaryote (Fungi), as well as seven Viruses (Fig. 1D). In terms of microbiome functions, the predominant COG categories were “G-carbohydrate transport and metabolism”, “J–Translation, ribosomal structure and biogenesis”, and “C–Energy production and conversion” (Fig. 1E, Fig. S6F). On the other hand, host human proteins were primarily associated with “O–Posttranslational modification, protein turnover, chaperones”, “R–General function prediction only”, and “Z–Cytoskeleton” (Fig. 1F). This underscores distinct functional profiles between microbial and human proteins within the human gut.

### Population-scale comparison on metaproteomics and metagenomics

To emphasize the unique insights gleaned from metaproteomics, we conducted an in-depth comparison between the metaproteomic data and the metagenomic data ^41^ of the GNHS cohort. Multiple levels, including human protein, metaprotein, COG, KO cluster, and taxonomy were employed to capture the microbial taxonomic and functional diversity within the GNHS cohort. Analysis was conducted on non-redundant samples collected from participants who had not received antibiotics (n = 1385) (Fig. 2A). Our findings showed that, as sample size increased, metaproteomic data reached maximum estimated richness with 333 individuals at the genus level and 302 individuals at the species level. For functional cluster KO, the metaproteome data achieved 99% of the maximum estimated richness with approximately 200 samples and reached the maximum estimated richness with 1,049 individuals (Fig. 2B, Fig. S7A). Additionally, metaproteomic data covered over 99% of the estimated maximum richness at the human protein, metaprotein, and COG cluster levels within 500 samples (Fig. S7A). In contrast, the metagenomic data of the GNHS cohort could only reached 85% of the saturation level at the taxonomic genus level, 81% at the species level, 93% at the KO functional cluster level, and 97% at the COG level with 1,219 individuals (Fig. 2B, Fig. S7B). Previous population-based metagenomic studies have shown that the observed richness is lower than the estimated richness even with sample sizes exceeding 8,000 ^40^. We also observed that the diversity of microbial taxa and their functions predicted by metagenomics far exceeds those detected by metaproteomics. Therefore, we infer that metagenomics more accurately represents the entire microbial composition and the spectrum of its genetic functions. Conversely, metaproteomics is more adept at identifying actively living microbes and their actively expressed functions. Despite abundant estimations in gut microbiome composition diversity among individuals, the diversity of expressed functions appears to be relatively conservative and less diverse. Consequently, metaproteomic studies can draw robust conclusions with fewer samples.

**Figure 2.**
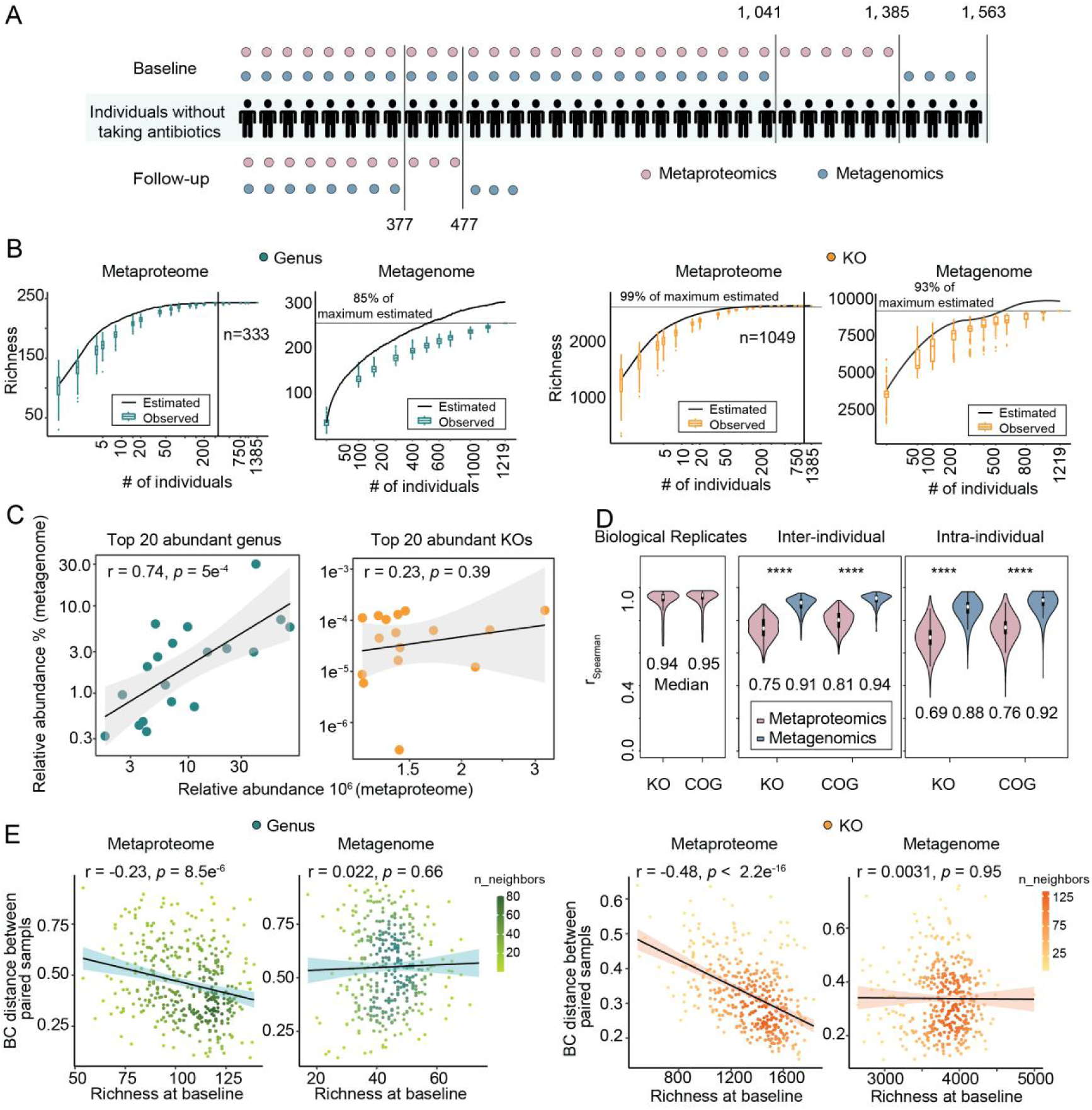
Comparison of metaproteomic and metagenomic data of paired samples. (A)The GNHS cohort collected fecal samples during baseline and follow-up, which were then subjected to metaproteomic or metagenomic analysis. (B) Genus or KO accumulation curves of metaproteomic and metagenomic data. The x-axes represent the cumulative sample number, and the y-axes mean the total richness of each feature. The black curves show the estimated richness; the box plots report the measured richness. The maximum measured richness and its proportion to the maximum estimated richness are indicated by a horizontal line. The vertical lines represent the minimum sample numbers required for full representation of GNHS cohort diversity. (C) Quantitative correlation analysis between the top 20 abundant genera or KOs in metaproteomics and their corresponding values in metagenomics. The x-axes show the relative abundance measured by metaproteomic data, and the y-axes are the quantitative value in metagenomic data. The correlation was calculated using Spearman. (D) The correlation of microbial functions between different samples or two samples collected from the same individual. The statistical analysis was performed with Welch’s t-test. *** *p* value < 0.001. (E) The temporal stability of the microbial genus or KO functions was associated with corresponding richness at the stage of baseline in metaproteomic data and metagenomic data. The correlation was calculated using Spearman.

Subsequently, we examined the correlation within and between metaproteomic and metagenomic datasets, focusing on paired metaproteomic and metagenomic data from 1,041 individuals with non-redundant sample collection (Fig. 2A). Apart from the human protein level, the BC distance data of microbial levels exhibited strong correlation with each other within each omic data (> 60% correlation) (Fig. S7C). We also observed statistically significant correlation between the two omic datasets. Notably, the correlation between the taxonomic datasets of the two omics was higher than that of the functional datasets (Spearman rho: 0.34 for the genus, 0.43 for species compared to 0.26 for COG, 0.25 for KO) (Fig. S7C), suggesting taxonomic data are more consistently aligned across omics than functional data. This was further supported by the distribution of abundant genera and KOs, where taxonomic correlation was typically higher. The most abundant taxa in two omics datasets were found to be consistent (Fig. S8 A-B). Among the top 20 genera identified in the metaproteomic data, 19 were also detected in the metagenomic data, exhibiting a strong correlation (r_Spearman_ = 0.74, *p* = 5e-4) in their relative abundance (Fig. 2C). A similar pattern was observed at the species level (r_Spearman_ = 0.72, *p* = 0.0052) (Fig. S8C). However, at the KO and COG functional levels, the highly abundant functions are distinct between the two omics datasets (Fig. S8 D-E). Although the highly abundant functions in the metaproteomic data were also detected in the metagenomic data, the quantitative results showed no statistically significant correlation between the two omics datasets (Fig. 2C, Fig. S8F). The observed differences in functional profiles between the two omics data might arise from the inherent measurement focuses: metagenomics quantifies gene copy numbers, whereas metaproteomics assesses protein expression ^42,43^.

Population-based metagenomic studies reported conserved functional category variations between individuals ^4,40,44^. Given the quantitative functional discrepancies between metaproteomics and metagenomics, we investigated whether metaproteomics could reflect the functional variation within the population. Analyzing the feces of 1,041 subjects with both metagenomic and metaproteomic data, we observed more pronounced variability in metaproteomics KO and COG categories compared to that in metagenomics (Fig. 2D). Additionally, our analysis revealed that variation in functional categories between baseline and follow-up samples were higher in the metaproteome data than in metagenomic data across 377 individuals (Fig. 2D). Thus, we inferred that metaproteomics may offer a more sensitive approach to detecting functional differences. Finally, we examined the long-term stability of the gut microbiome within individuals. Both metaproteomics and metagenomics data showed that individuals displayed stable gut microbiome compositions and functions for over 1.4–5.5 years (Fig. S9 A–B), consistent with a previous metagenomic study ^45^. It has been reported that the stability of the microbiome over time inversely correlates with its biodiversity at the initial assessment ^45^. In our data, the metagenomic analysis showed a slight negative correlation (r_Spearman_ = −0.1) between individual microbiome variation and the baseline taxonomic species richness. No significant correlation could be obtained at metagenomic taxonomic genus, functional KO, or COG levels (Fig. 2E, Fig. S9C). In contrast, metaproteomic analysis revealed significantly negative correlations across all microbial taxonomic and functional levels. Notably, the correlation was stronger at the microbial function level than the microbial taxonomy level (r_Spearman_ = −0.23 for genus vs. r_Spearman_ = −0.48 for KO function), suggesting that functional diversity, as captured by metaproteomics, provides a more reliable measure of microbiota stability over time.

In summary, our study highlights the superior capabilities of metaproteomics in revealing the functional dynamics of the microbiota. Through the lens of metaproteomics, we were able to capture a comprehensive view of microbial functional biodiversity within a population of fewer than 500 individuals, identify significant functional variations, and gain deeper insights into the temporal dynamics of microbiome functions.

### Core microbial functions and the core species expressing these functions

Core features are believed to play crucial roles in maintaining the health of the gut ecosystem. To identify core features within the GNHS cohort, we conducted a screening for attributes shared across individuals. Features present in more than 80% of both inter-individual (non-redundant samples collected from 1,041 individuals) and intra-individual samples (samples collected at the baseline and follow-up time points from 377 individuals) were categorized as “core”. In the metaproteomic data, 262 human proteins, 353 metaproteins, 9 phylum, 56 genera, 62 species, 797 KOs, and 947 COGs emerged as core features (Fig. S10A, C–E; Table S2). The metagenomic data revealed 4 phyla, 19 genera, 23 species, 3,227 KOs, and 2,773 COGs as core features (Fig. S10B, F–H; Table S3).

Our primary focus is on the core microbial functionalities present within the population and how these functions are distributed among the core microbes. To pinpoint core microbial functionalities, we characterized pathways significantly enriched in core KOs as “core pathways”, using a criterion of a Benjamini & Hochberg (BH) adjusted p-value ≤ 0.05 and a fold enrichment value ≥ 2. Additionally, we mapped the uniquely expressed metaproteins of each core species and identified the pathways that were significantly expressed by them (with a BH adjusted p-value ≤ 0.05 and fold enrichment ≥ 2). This dual analysis allows us to discern which microbial pathways are likely to be expressed and the core microbes responsible for their expression. Our findings revealed 42 core pathways enriched in metaproteomics and 64 in metagenomics, with 39 pathways common to both datasets (Table S4). Notably, core pathways in metaproteomics showed higher fold enrichment values than those in metagenomics (Fig. S11), indicating a greater likelihood of these pathways being actively expressed. We classified pathways with greater fold enrichment in metaproteomics compared to metagenomics as “highly expressed pathways” and pathways with lower fold enrichment in metaproteomics as “moderately expressed pathways”. We found that highly expressed pathways mainly involve “housekeeping” functions essential for microbial survival, such as translation, energy metabolism, carbohydrate metabolism, and amino acid metabolism. These functional pathways, significantly enriched within the core microbes and detected in at least 15 core species (Fig. S11, Table S5), exemplify functionalities commonly expressed across the microbial community. This aligns with findings from population-based metatranscriptomic studies, suggesting these housekeeping functions are readily transcribed ^46^. Furthermore, the enriched pathways occurred in a limited number of core species, involving functions like glycan degradation, streptomycin biosynthesis, drug metabolism related to xenobiotic degradation, as well as taurine and hypotaurine metabolism. Despite being expressed by only a few core microbes, these pathways still exhibited relatively high expression in the population (Fig. S11, Table S5). In contrast, the moderately enriched core pathways within the population are mainly associated with environmental stress response, such as cellular community, cell motility, replication and repair, and membrane transport. These functions, enriched in fewer than three core microbes, exhibit low expression levels across the population (Fig. S11, Table S5). This distinction between highly and moderately expressed pathways provides insight into the functional stability and adaptability of the core microbes within diverse environments.

The metabolism-related pathway constituted the largest proportion of highly expressed group in gut microbe community, underscoring the microbial community’s active metabolic function. Short-chain fatty acids (SCFAs) and vitamin Bs act as an energy source for intestinal cells and regulate the activity of the host’s immune ^47–49^. According to our data, a total of 33 microbes were found to express functions related to the metabolism of SCFAs or vitamin B, with 23 of these microbes previously reported to have the capability of secreting these beneficial metabolites (Fig. 3A, Table S6). We also found core microbes that contribute to the production of SCFA or vitamin B which have not been reported. Notably, *Frisingicoccus caecimuris* demonstrated a significant capability for propanoic acid production, evidenced by a high fold enrichment value of 42 in the propanoate metabolism pathway. *Clostridium fessum* showed notable activity in butanoate metabolism with a fold enrichment of 18, while *Blautia hansenii* was involved in vitamin B9 metabolism, presenting a fold enrichment value of 22 (Fig. 3A and Table S6). To verify our findings, we measured the butyric acid content in feces from 494 individuals. Our analysis revealed a positive correlation between the total relative abundance of 19 specific bacteria and the level of butyrate in the fecal samples, suggesting the potential contributions of the microbiome community to butyrate production (Fig. 3 B-C, r_Spearman_ = 0.25, *p* = 2.7e^-8^).

**Figure 3.**
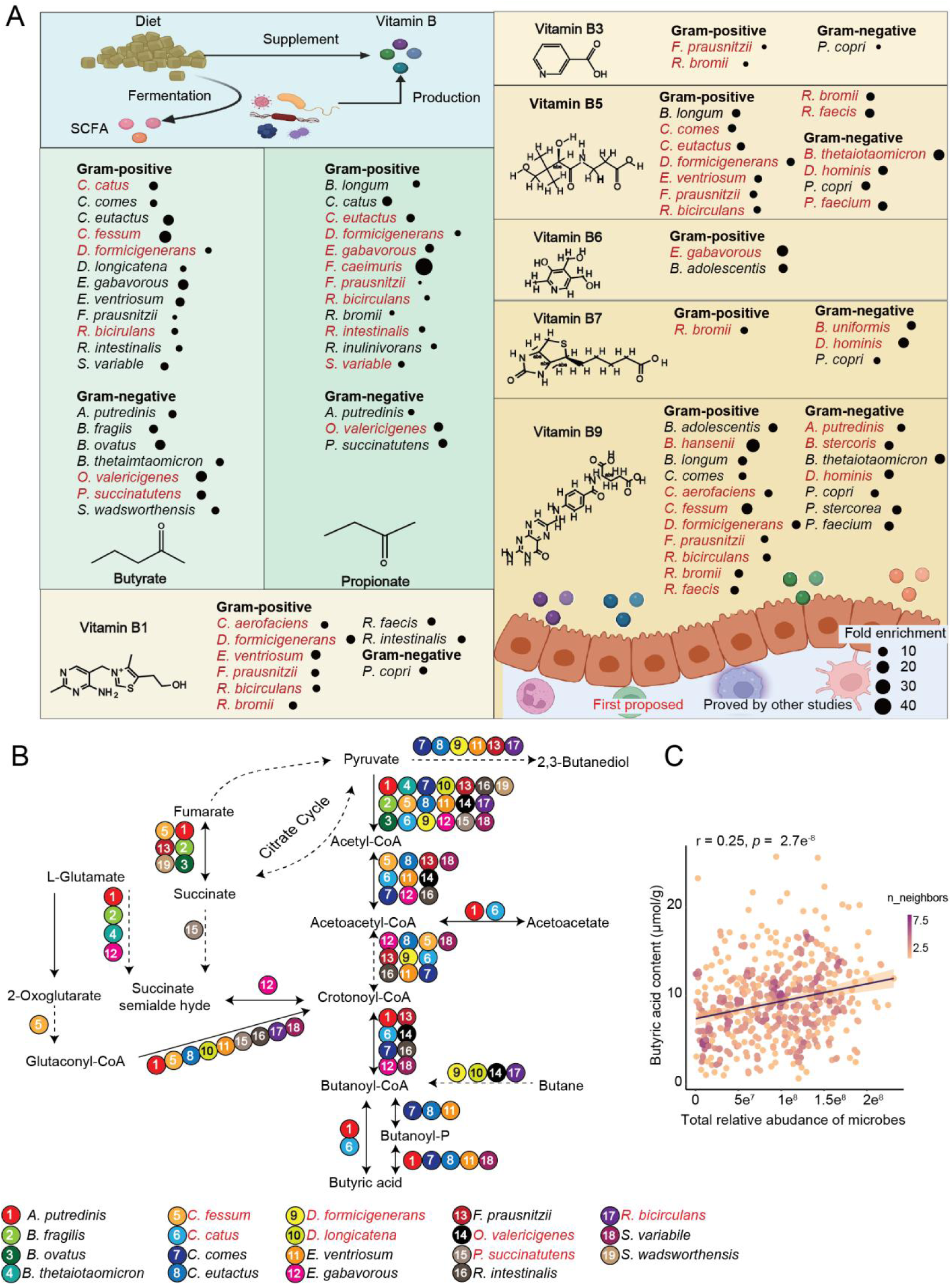
Metaproteomic core species involved in the metabolism of SCFA and vitamin Bs. **(A)** Summary of metaproteomic core species enriched with SCFAs (butyrate and propionate), as well as vitamin Bs (B1, B3, B5, B6, B7, and B9) metabolisms. The dots represent the fold enrichment value of pathways in each species. The bacteria denoted in black font are consistent with the reports, while those in red were firstly found in this study. (B) Metaproteomic core species highly expressing butyric acid metabolism were summarized, with bacteria capable of detecting the involved enzyme labeled next to the reaction. The bacteria denoted in black font are consistent with reported butyric acid-producing species, while those in red were firstly found in this study. (C) The fecal butyric acid content was found to be correlated with the total abundance of core microbes highly expressing butyric acid metabolic pathways identified in metaproteomics. The correlation was calculated using Spearman.

In conclusion, we have delineated the highly expressed biological pathways and identified the core species responsible for these expressions. We also identified potential core microbes capable of producing SCFAs or Vitamin Bs, emphasize the crucial role of metaproteomic analysis in accurately identifying microbial contributors to key metabolic processes.

### Associations between phenotypes and metaproteomic data

To explore the relationship between phenotypic variation and the diversity of gut microbiome among individuals, we performed permutational multivariate analysis of variance (PERMANOVA) to assess the association between 49 phenotypes (excluding antibiotic use) and BC dissimilarities at various levels, including human protein, microbial metaprotein, KO, COG, genus, and species levels. Five metadata parameters, including type 2 diabetes medicine (T2D medicine), blood triglyceride (TG) level, Bristol scale, T2D, and tea consumption, were found to exhibit significant association with BC distance variation at all levels (Fig. S12, adjusted p-value ≤ 0.1, Table S7). Among these metadata, the most significant contributor was T2D medicine, followed by TG level, Bristol stool scale, T2D, and tea consumption. These results align with findings from population-based metagenomic studies conducted in various countries and regions, reinforcing the impact of these phenotypes on the gut microbiome’s composition and function ^4,40,50–52^. We next explored the specific associations between phenotypes and the metaproteomic data using general linear model (GLM) method. A total of 10,714 associations were identified between metaproteomic data and 39 phenotypes (Table S8). Still, T2D medicine, Bristol scale, T2D, TG, and tea consumption emerged as the primary phenotypes correlated with metaproteomic data across various levels, including the levels of protein (metaprotein and human protein), microbiota function (COG and KO), and taxonomy (genus and species) (BH adjusted *p* ≤ 0.05, Fig. 4A).

**Figure 4.**
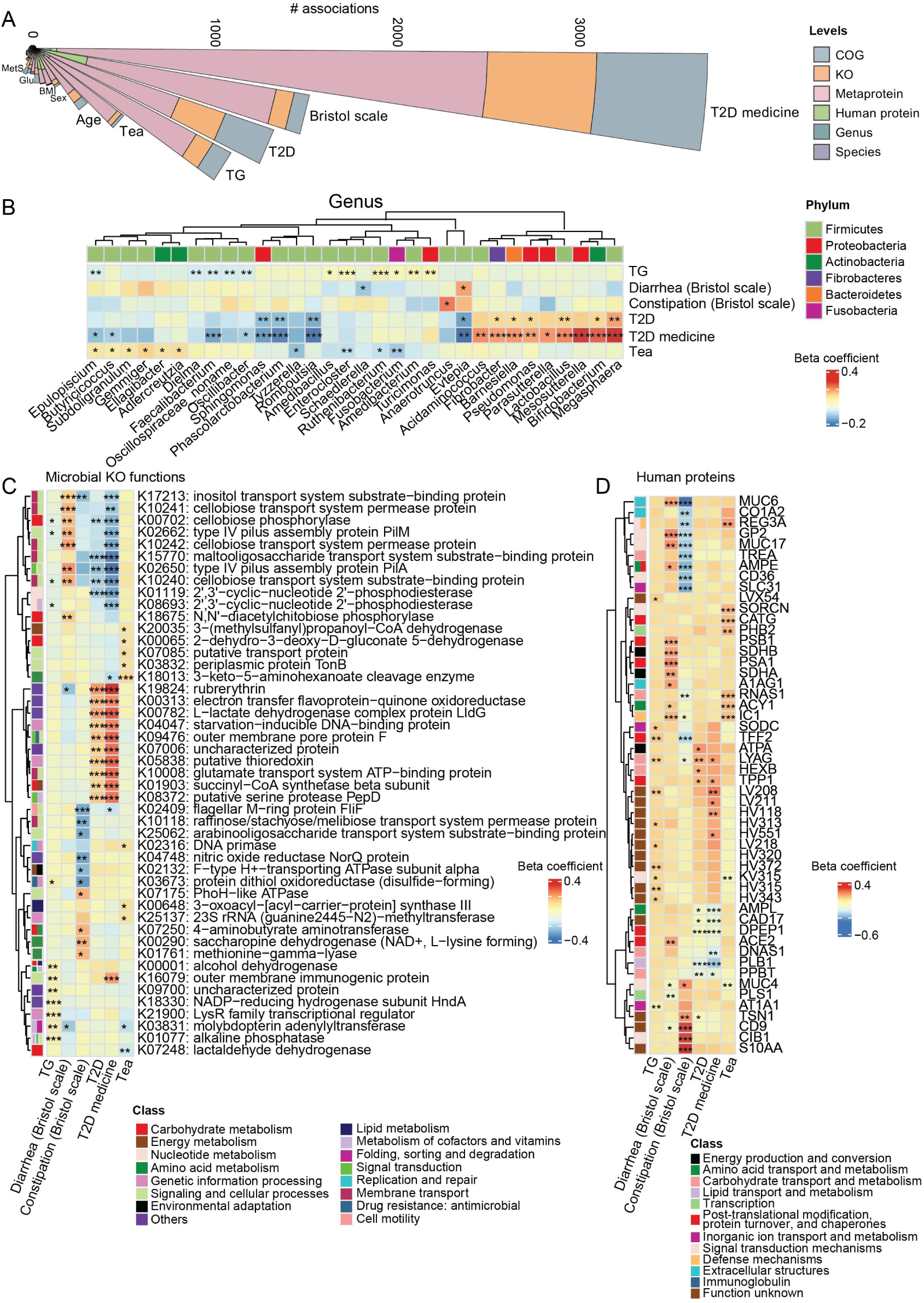
Associations between phenotype and metaproteomic features. (A) Summary of the associations between metaproteomic characteristics and phenotypes. The bars in the graph represent different phenotypes and are sorted according to the number of associations for each level. (B) Metaproteomic genera with the strongest overall correlations with five covariates – Bristol scale, T2D, T2D medicine, TG, and tea. The beta coefficients of each feature and phenotype were plotted along with their corresponding BH-adjusted p values, where * represents p < = 0.05, ** < = 0.01, *** < = 0.001. (C) Metaproteomic microbial KO functions with the strongest overall correlations with covariates. (D) Metaproteomic human proteins with the strongest overall correlations with covariates.

Some of the associations at the taxon level have been previously reported. For instance, the genus *Anaerotruncus* was positively correlated with constipation. *Megasphaera* showed a positive correlation with T2D ^53^. *Evtepia* and *Romboustsia* ^54^ exhibited a negative correlation with T2D. Tea drinkers displayed increased *Gemmiger* ^55^ and decreased *Tyzzerella* levels ^56^. In addition, at the function levels, we uncovered protein functions of microbes and the host that are significantly associated with these metadata for the first time. At the microbial KO level, enzymes related to carbohydrate metabolism and energy metabolism were significantly increased in diarrhea and significantly decreased in constipation. Amino acid metabolism functions were upregulated in constipated individuals. Diabetic patients showed decreased expression of carbohydrate metabolism and energy metabolism proteins, while the levels of rubrerythrin, electron transfer flavoprotein-ubiquinone oxidoreductase, and lactate dehydrogenase were elevated. At the human protein level, mucin proteins MUC6 and MUC17, proteasome PSA1 and PSB1, as well as succinate dehydrogenase enzymes SDHA and SDHB, were increased in diarrhea individuals. In individuals with constipation, the abundance of mucin proteins as well as carbohydrate degradation related trehalase TREA and alpha-amylase-associated protein SLC3 were decreased. (Fig. 4D).

Aging is the root cause of numerous diseases and functional disorders. The connection between aging and the gut microbiome has become increasingly emphasized ^57,58^. Beyond the five phenotypes previously discussed, our data also revealed age-related gut microbes and their functions. At the microbial functional level, there were 58 microbial KOs associated with aging (Table S7). Among these, glycine dehydrogenase, beta-glucuronidase, and proline racemase exhibited the strongest negative correlation with aging, indicating a potential decline in the overall metabolic activity of the gut microbiome with increasing age (Fig. S13A). Conversely, outer membrane proteins of Gram-negative bacteria, such as ompF, ompC, and lamB, showed the strongest positive correlation with aging. Outer membrane proteins play a crucial role in pathogens, aiding bacteria in evading host defense mechanisms, thereby increasing their virulence^59^. Therefore, the increase in outer membrane proteins might suggest that with age, the activity or potential pathogenicity of certain pathogenic strains in the gut microbiome could increase, potentially impacting the host’s health. Taxonomically, four genera were associated with aging, including negative associated genus *Phascolarctobacterium* and positive associated genera *Klebsiella*, *Dialister*, and *Veillonella* (Fig. S13B). By connecting these microbial genera with their unique functional expressions, we observed specific changes in the functionality of each microbe. Functions related to energy metabolism, lipid metabolism, and nucleotide metabolism of *Dialister* showed positive correlation with aging. Similarly, energy metabolism functions of *Veillonella* were also positively associated with aging. Additionally, the outer membrane porins primarily expressed by *Klebsiella* were positively correlated with aging, indicating an increased pathogenicity as age advanced (Fig. S13C).

The correlation between metaproteomic data and phenotypes deepen our understanding of the changes in the gut microbiome and their functions under different phenotypic conditions, thus offering new perspectives in disease prevention and health management.

### T2D related microbes and functions identified through metaproteomics

T2D has been proven to be related to gut microbiota composition, but the specific functions of microbiota associated with T2D remain incompletely understood. In this study, we analyzed non-redundant samples from 1153 non-T2D and 223 T2D individuals within the GNHS cohort to investigate T2D-associated metaproteomic features (Fig. 5A). A total of 1,381 features including 785 metaproteins, 23 human proteins, 268 KOs, 279 COGs, 14 genera, and 12 species were identified significantly associated with T2D (BH adjusted p ≤ 0.05, Table S8). Among these, 335 metaproteins were uniquely expressed by 36 microbial species (Table S9). Consequently, a total of 40 species were identified to be correlated with T2D when considering the combined results of species related to T2D and those corresponding to T2D-related metaproteins. These findings are consistent with previous reports, where 25 species were identified as related to T2D using 16S or metagenomic sequencing (Table S9). Our data further elucidate the significantly expressed metaproteins of these bacteria, with over half of the species-uniquely-expressed metaproteins (176 out of 335) involved in carbohydrate, amino acid, lipid, and nucleotide metabolism, as well as energy production. This highlights the critical regulatory roles of these functions in T2D ^51,60–62^ (Fig. S14). Specifically, the 40 species mainly form 4 distinct microbial community clusters based on their interrelationships (Fig. S14, Fig. S15A). Cluster 1 included 6 species from the *Bifidobacterium* genus, with their abundance or protein expressions positively correlated with T2D. Specifically, these microbial proteins were mainly associated with amino acid transport and carbohydrate metabolism. Cluster 2 including 14 species, with 11 of them belonging to Eubacteriales. These bacteria mainly expressed carbohydrate metabolism related functions that were negatively correlated with T2D. The Cluster 3 was composed of *D. hominis* and *A. histamini*, both belonging to the family Veillonellaceae. They were positively correlated with T2D. Their proteins mainly participated in energy metabolism and amino acid metabolism. The last cluster consisted of two species belonging to the *Megasphaera* genus, positively correlated with T2D. The microbial proteins expressed in this cluster were mainly related to energy metabolism and lipid metabolism (Fig. S15A). To validate the T2D-related features identified in the GNHS cohort, we included an independent cohort, referred to as the FH cohort, consisting of 82 non-T2D individuals and 21 T2D individuals. Through this validation process, we confirmed the association of 45 metaproteins, 1 human protein, 27 KOs, 18 COGs, and 3 species associated with T2D (consistent orientation of associations and BH adjusted *p* ≤ 0.05, Table S10). Notably, *Megasphaera elsdenii* and 20 of its expressed metaproteins, predominantly involved in energy production and lipid metabolism, were confirmed to be associated with T2D in the FH cohort as well. These consistent associations suggested a potential significant role for *M. elsdenii* in the development of the disease (Fig. S15B).

**Figure 5.**
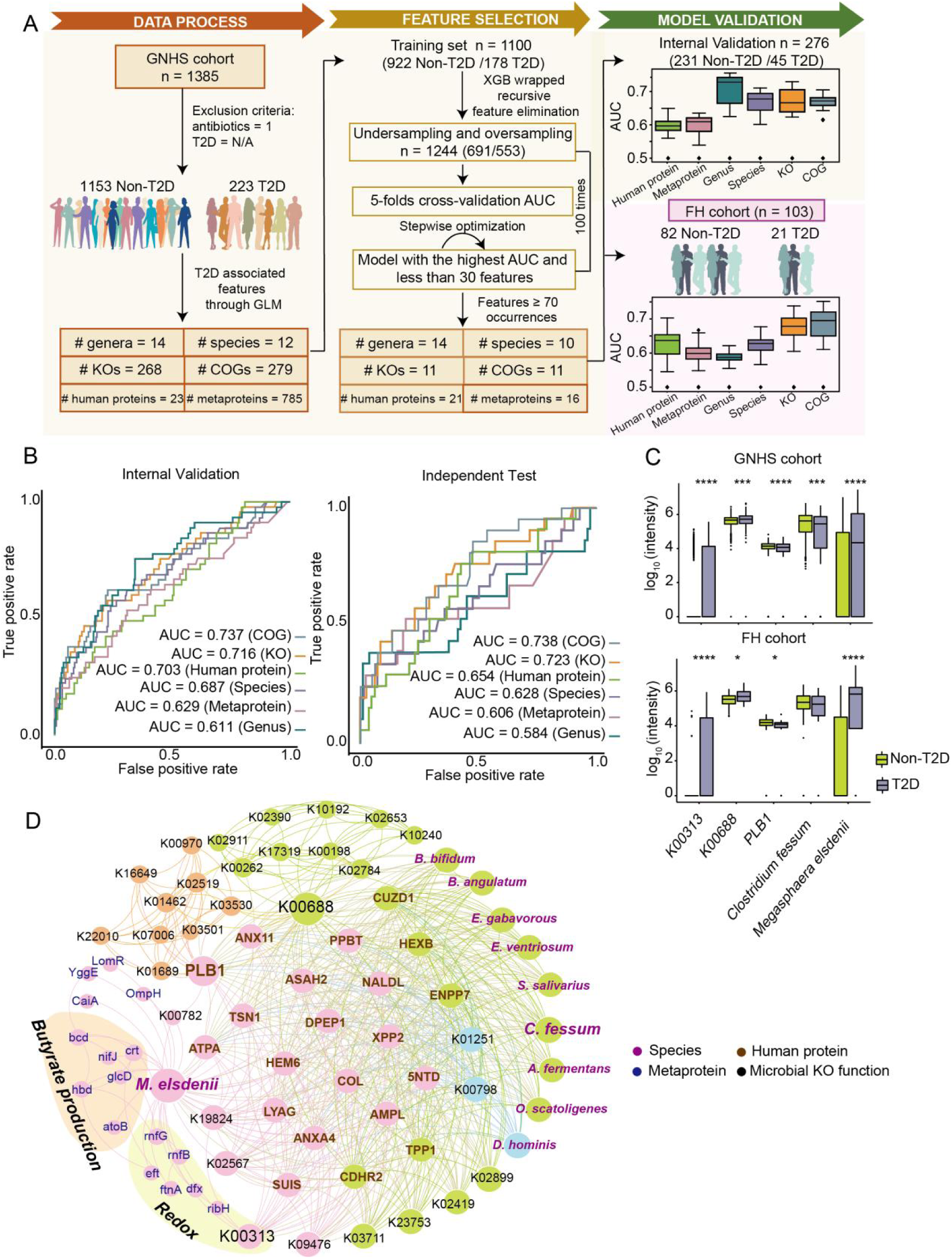
Metaproteomic functional insights of T2D. (A) The machine learning workflow for diabetes classification based on metaproteomic data includes data processing, feature selection, and model validation. Features with a q-value < 0.05 are filtered using GLM and used as input for diabetes-related features. Among the 100 machine learning models, features that occur more than 70 times are further selected, and their performance is evaluated in both the internal validation test and the independent test set at each level. (B) The receiver operating characteristic (ROC) curves showcasing the best performance for each level in the internal validation cohort and the independent FH cohort. (C) The log_10_ transformed intensity of the five potential T2D biomarkers in the GNHS cohort and FH cohort. Wilcoxon rank-sum test was used for the differential analyses. * *p* value < 0.05, *** *p* value <0.001, **** *p* value < 0.0001. (D) Networks of the five potential T2D biomarkers and their associated T2D-related features in GNHS cohort. Links between metaproteins and species, as well as KOs and species, were based on their common peptides, while the co-expressed linkages between intra-levels and inter-levels were calculated by the Spearman correlation (BH adjusted p-value < 0.05, | r | ≥ 0.3 for inter-levels, | r | ≥ 0.8 for intra-levels).

Next, we explored the performance of metaproteomic data in classifying T2D patients from those without disease. To achieve this, we utilized the T2D-associated features identified through GLM analysis in the GNHS cohort as potential features (Fig. 5A). These features were further refined using the eXtreme Gradient Boosting (XGBoost) algorithm with wrapped recursive feature elimination. Using this approach, we identified 16 metaproteins, 21 human proteins, 11 KOs, 11 COGs, 10 species, and 14 genera as the most robust features for machine-learning models at each level (Table S11, Fig. 5A, and Methods). The performance of these features was evaluated by an internal validation and the independent FH test cohort at each level. Notably, the microbial functional COG and KO levels exhibited strong performance in both the internal and test cohorts (Fig. 5B). The best receiver operating characteristic (ROC) area under the curves (AUCs) reached 0.738 and 0.723 at COG and KO levels in FH cohort, respectively (Fig. 5B). We explored the feasibility of using metagenomic data for T2D classification using the same approach (Fig. S16B). At the species level, the best AUC of the machine learning model achieved 0.62 in the independent validation set, comparable with that of metaproteomics. However, when evaluating microbial functional KO levels, metagenomics only achieved an AUC of 0.5 much lower than that of metaproteomics (Fig. S16C). These findings highlight the advantage of quantitative characterization of microbial expressed functions by metaproteomics in T2D classification. (Table S12, S13)

Combining the outcomes of GLM analysis and machine learning, we carried out an extensive screening to identify the most conclusive features associated with T2D. These features exhibited significant correlation with T2D in both the GNHS and FH cohorts. We identified five pivotal features, consisting of two microbial functions—electron transfer flavoprotein-quinone oxidoreductase fixC (K00313) and glycogen phosphorylase (K00688), two bacterial genera—*Clostridium fessum* and *M. elsdenii*, and one human protein—phospholipase B1 (PLB1) (Fig. 5C). *M. elsdenii,* K00313, and K00688 were elevated in T2D patients, and their relationships with diabetes have been reported ^63–65^. While PLB1 and *Clostridium fessum* were first found to be decreased in T2D patients. A co-occurrence network was developed to illustrate the regulatory roles of machine learning and FH cohort validated T2D-related features (Fig. 5D). Notably, *M. elsdenii* emerged as a hub microbe with the highest connectivity in the network. The network specifically highlights up-regulated proteins in the *M. elsdenii,* primarily concentrated in butyrate production through lactate utilization and redox pathways. This emphasizes the critical role of *M. elsdenii* and its functions in T2D. (Fig. 5D).

### *M. elsdenii* safeguards against diabetes by producing butyric acid

Next, we directed our analysis towards *M. elsdenii* to explore its potential role in the development of T2D. Supported by prior research ^66,67^, our findings showed that *M. elsdenii* utilizes lactate as the energy source to produce butyric acid. Given that butyric acid has been recognized to alleviate diabetes symptoms ^68,69^, the increased abundance of *M. elsdenii* in T2D patients might initially appear paradoxical. However, our investigation also uncovered significant positive correlations between *M. elsdenii* and its proteins with antidiabetic drug (Fig. S18; Table S9), a notable observation considering about 57% of diabetic patients were under medication treatment. This led us to speculate that the observed positive correlation between T2D and *M. elsdenii* could primarily stem from the medications commonly prescribed to these patients. Analyzing data from three distinct groups—non-diabetic individuals, untreated diabetics, and diabetics on medication, we noted an increasing trend in *M. elsdenii* levels among patients undergoing antidiabetic treatment (Fig. 6A). Corroborating our hypothesis, previous studies have documented elevated levels of *M. elsdenii* in patients receiving antidiabetic treatment, including metformin and acarbose ^70,71^. To confirm the direct effect of antidiabetic medication on microbial growth, we conducted *in vitro* experiments by introducing common anti-diabetic drugs, namely metformin, acarbose, and glimepiride, to a pure culture of *M. elsdenii.* The outcomes, measured through both the OD600 values and bacterial cell counts, verified that acarbose and metformin promoted the growth of *M. elsdenii*, with the exception of glimepiride (Fig. 6B).

**Figure 6.**
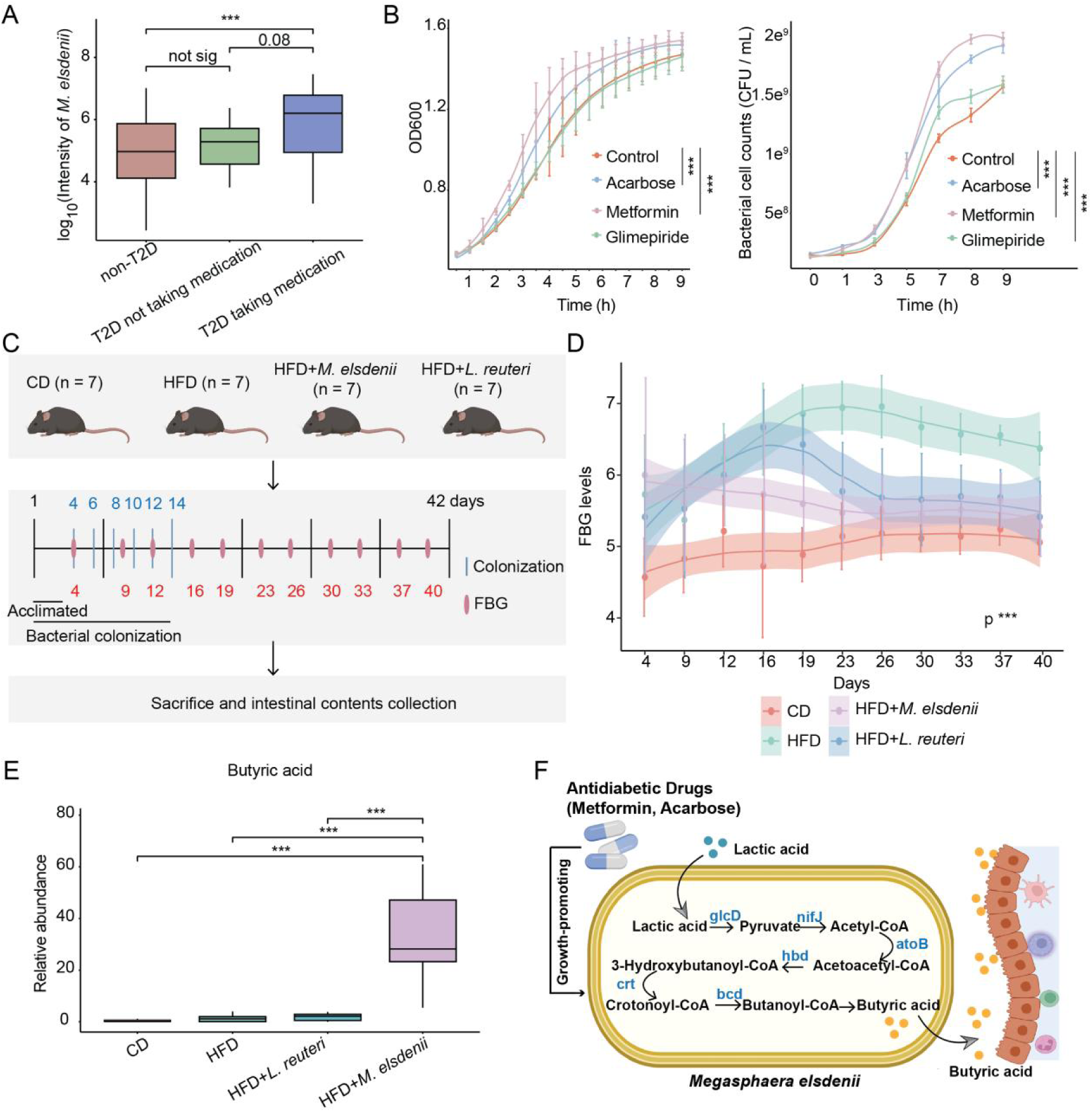
Biofunction of *M. elsdenii*. (A) The log_10_ transformed intensity of M. elsdenii in non-T2D, T2D not taking medication, and T2D taking medication groups. Wilcoxon rank-sum test was used for the differential analyses. *** *p* value <0.001. (B) Growth curves of *M. elsdenii* in the presence or absence of exogenous drugs assessed by measuring the OD600 value of the bacterial culture and counting the number of live bacterial cells, respectively. Two-way ANOVA with repeated measurement was applied to calculate the difference between control and drug-treatment groups. *** *p* value <0.001. (C) Experimental design of germ-free mouse study. Mice were fed with either a CD or a HFD for 42 days. To investigate the impact of *M. elsdenii* and *L. reuteri* on blood glucose, mice fed with HFD were monocolonized with these two strains for the initial 14 days of the study. Throughout the experiment, FBG levels were monitored twice weekly. After the study period, the intestinal contents of the mice were collected to analyze the levels of SCFAs in the gut. (D) The FBG levels of each mice group during administration. Two-way ANOVA with repeated measurement was applied to calculate the difference between the HFD and HFD+FMT(*M. elsdenii*) groups. *** *p* value <0.001. (E) The butyric acid levels in the intestinal contents of each mice group. Wilcoxon rank-sum test was used for the differential analyses. *** *p* value <0.001. (F) The pathogenesis of *M. Eisdenii* in inducing diabetes based on metaproteomic and experimental evidence.

The increase of *M. elsdenii* through drug treatment might suggest its glucose-lowering effect on T2D. To explore this possibility, we conducted an experiment with gnotobiotic mice on a high-fat diet (HFD), treating them with *M. elsdenii* and a control bacterium, *Lactobacillus reuteri*, known for its glucose-lowering effects in mice ^72,73^. The mice were subjected to an HFD for 42 days, with the first 14 days including bacterial colonization. We monitored their fasting blood glucose (FBG) levels throughout the study (Fig. 6C). Compared to the control group on a chow diet (CD), mice on the HFD exhibited significantly higher FBG levels. However, the group treated with *L. reuteri* showed a consistent reduction in FBG levels compared to those on the HFD after 26 days (Fig. 6D). Similarly, mice fed with *M. elsdenii* showed a notable and consistent decrease in FBG levels stating from day 16, as compared to HFD-fed mice (Fig. 6D), demonstrating its glucose-lowering impact on T2D. Our metaproteomic data revealed an increase in proteins related to butyric acid production in *M. elsdenii*. To investigate the mechanism underlying blood glucose levels induced by *M. elsdenii*, we examined the intestinal contents of mice after 42 days to assess the level of butyric acid. We found that HFD mice fed with *M. elsdenii* exhibited significantly higher butyrate level than the other groups (Fig. 6E), suggesting a possible mechanism that *M. elsdenii* contributes to glucose regulation in T2D.

Therefore, drawing from the metaproteomic data, *in vitro* culture, and mouse experiments, we found that *M. elsdenii* responds to the stimulation induced by common antidiabetic drugs like acarbose and metformin. This stimulation enhanced the production of butyric acid, which subsequently reduces high blood glucose levels and alleviates diabetes conditions (Fig. 6F).

## Discussion

In this project, we have developed a metaproteomics analysis pipeline tailored for large-scale population samples. This pipeline demonstrated high-throughput capability and reproducibility. We applied this method in a regional cohort from China, which included over a thousand individuals. Notably, this represents the inaugural application of metaproteomics on such a sizable cohort. The critical bottleneck in metaproteomics analysis lies in aligning microbial proteins with mass spectrometry spectra. Given the limited understanding of the full spectrum of gut microorganisms, there is a lack of a comprehensive database to underpin this process. To overcome this obstacle, numerous studies have utilized paired metagenomes to construct protein databases. This approach facilitates the analysis of metaproteomics data and aids in identifying microbial proteins most pertinent to the metagenomic context. For example, open-access resources like the Integrated Gene Catalog (IGC) ^74^, derived from integrated metagenomic sequencing data, and the Unified Human Gastrointestinal Protein Database (UHGP) ^75^, built on sequenced microbial genomes, are increasingly being integrated into metaproteomic analyses. However, relying on metagenomic sequencing to generate a reference database adds extra burden to metaproteomic analysis. Here, we utilized an open-source database instead of paired metagenomes to align the microbial protein sequences with mass spectrometry data. Additionally, the IGC database, derived from metagenomic sequences of 1,070 individuals, notably includes a significant representation from China. This makes it particularly suitable for exploring Chinese cohorts. As a result, we have chosen the IGC database as the foundation of our reference database.

Through metaproteomic analysis, we identified 1,277 unique human proteins and 86,876 microbial proteins from 1,967 fecal samples of 1,399 individuals. One of the key advantages of metaproteomics over metagenomics is its capability to detect host proteins. However, accurately determining the origin of these host proteins, whether they arise from shed epithelial cells, host-secreted proteins, or genuine interactions with microbes, is crucial as it directly impacts the scientific significance of our protein detection findings. To address this, we initially employed a differential centrifugation method to remove shed epithelial cells and food residues from the fecal samples before enriching the microbial fraction for metaproteomic analysis. This step effectively eliminated the majority of host proteins derived from shed epithelial cells. Furthermore, we performed multiple rounds of differential centrifugation washing to ensure the comprehensive removal of impurities while verifying the presence of proteins. Across the 3 to 5 washing stages, we consistently detected these host proteins, indicating their stable association with microbial precipitates. Functional annotation analysis of the identified host proteins revealed their predominant involvement in food digestion enzymes, with some confirmed to interact with microbial proteins ^39^. Consequently, we believe that the identification of these human proteins provides valuable insights for future investigation into the complex interplay between the host and gut microbiota.

Metagenomics and metaproteomics are vital techniques for exploring microbial species and their proportions through gene or protein analysis. While comparisons between these methods have shown some consistency in classifying high-abundance microbial populations ^20,75^, there is a lack of research on the advantages of metaproteomics over metagenomics in population studies and disease applications, primarily due to limited sample sizes. In our study, leveraging data from thousands of samples, we found that metaproteomic data tends to be more conservative compared to metagenomic data in discerning population diversity. Metaproteomics can capture the diversity of microbial classification within a population with around 350 samples and detect all microbial functional clusters expressed by the microbes with approximately 1000 samples. Our findings, based on a population of middle-aged and elderly individuals in China, suggest that metaproteomics offers advantages in encompassing the diversity of microbial populations in human populations, particularly in capturing active microbial functions in the gut. Considerations must be given to the limitations of metaproteomics, including the sensitivity of mass spectrometry, which can impact protein detection depth, and the prevalence of highly abundant microbial proteins in the results. While the conclusions regarding high-abundance microbial compositions in metagenomics and metaproteomics are generally congruent, the functional expression of microbial populations exhibits little correlation between the two omics approaches. Across populations or in samples collected repeatedly from the same individual over years, microbial protein expression in metaproteomics displays greater variability compared to gene content detected in metagenomics. Large-scale metagenomic studies have shown that while there is substantial variability in microbial composition among populations, the abundance of gene functions remains relatively consistent ^40,44^. The increased functional variability in metaproteomics relative to metagenomics may stem from differences between gene copy number and protein expression. Our study also highlights the benefits of metaproteomics for longitudinal studies of microbial populations within individuals over several years. The correlation between the biodiversity and the stability of microbial composition and function within an individual over time is stronger in metaproteomic data compared to metagenomics, making it a valuable tool for investigating the dynamics of microbial communities in human cohorts. Overall, while metagenomics and metaproteomics provide complementary information, metaproteomics offers unique insights into microbial diversity, functional expression, and longitudinal stability within human populations. The combination of these omics’ approaches can enhance our understanding of the complex interactions between host and microbiota in health and disease contexts.

In the analysis of core microbial features in the GNHS cohort using metaproteomic and metagenomic data, we observed differences of the core microbial taxa at the genus level and core microbial functional clusters at the KO level between two omics data. Metaproteomics revealed a greater number of core microbial taxa at the genus level (56 in metaproteomics vs 19 in metagenomics) compared to metagenomics, while metagenomics identified fewer core microbial functional clusters at the KO level (797 in metaproteomics vs 3,227 in metagenomics). The core genera identified in the metaproteome mainly encompassed those found in the metagenome, and similarly, the core functions identified in the metagenome largely encompassed those found in the metaproteome. The discrepancy of the core genus between two omics may be attributed to differences in the annotation databases for microbial classification. Utilizing the open-source IGC protein sequence database for metaproteomics could lead to variations in taxonomic annotation compared to paired metagenomic data. However, despite these differences, abundant core genera such as *Prevotella*, *Faecalibacterium*, and *Bacteroides* remained consistent between the two omics data. These genera have been previously identified as core genus in worldwide population studies based on 16S rRNA gene sequencing and metagenomics ^5,76^, further emphasizing their significance in the gut ecosystem. When comparing microbial functions, the disparities between the core microbial functions identified by the two omics approaches may arise from whether genes are actively expressed as proteins. Beyond transcription and ribosomal proteins, the core-expressed microbial functional groups predominantly encompass metabolic processes, including sugar metabolism, amino acid metabolism, lipid metabolism, and energy production. These findings underscore the significance of intestinal microbes in modulating host metabolic pathways for overall environmental health. Enriching these core microbial functions into pathways revealed that widely expressed microbial functions in the population are mainly related to microbial survival, with significant functional redundancy among microbes. These findings are consistent with the results from a metatranscriptomics study ^46^. However, within the core functional pathways, certain functions such as glycan degradation, streptomycin biosynthesis, xenobiotic degradation, as well as taurine and hypotaurine metabolism, are expressed by only a limited number of core microbes. This suggests that these microbes and their functions may play a central and potentially irreplaceable role in the microbial community, indicating their importance in maintaining microbial community stability.

We delved deeper into the distinct microbial proteins expressed by each microbe, delineating the independent high-expression functions of each organism. This involved functional annotation and pathway enrichment analysis of these metabolic proteins, with a particular focus on the vital roles of SCFAs and vitamin Bs in microbial contributions to host health. We pinpointed 33 core species potentially involved in the metabolism of SCFAs and vitamin Bs. Remarkably, 23 of these species have been previously documented to harbor these metabolic capabilities, underscoring the utility of metaproteomic data in elucidating microbial contributions to industrial production. However, it’s imperative to acknowledge the constraints and limitations inherent in such analyses. Given the phenomenon of cross-feeding in the gut microbiota, where various microbes collaborate to fulfill common metabolic pathways ^77^, pinpointing specific microbes solely based on pathway enrichment merely indicates their potential involvement in SCFA and vitamin B metabolism. Further research is imperative to determine whether these microbes can autonomously undertake SCFA or vitamin B metabolism, or if they rely on assistance from other microbes.

We also investigated the associations between metaproteomic data and 49 phenotypes. We found that T2D medicine, TG level, Bristol scale, T2D, and tea consumption exhibit the strongest associations with microbial features. This finding is consistent with the observations from other population-based metagenomic studies ^50,78,79^. We identified the microbes and their functions most closely associated with the five phenotypes and aging. Noteworthy correlations include *Anaerotruncus* with constipation, *Megasphaera* with T2D, *Klebsiella* with aging, and specific changes in microbial functions associated with these phenotypes.

Finally, our study centered on examining the microbiota and their functions associated with T2D. While previous metagenomic research has identified various microbiota linked to T2D ^80–83^, this study aimed to uncover the specific functional contributions of these microbiota to T2D. Our findings revealed that *Bifidobacterium* may influence T2D through amino acid transport and carbohydrate metabolism, while *Megasphaera* could contribute to T2D via energy and lipid metabolism. By employing a combination of GLM and XGBoost machine learning algorithms, we pinpointed five stable microbiota features consistently associated with T2D in both the GNHS and independent FH cohorts. Remarkably, we noted a noteworthy correlation between *M. elsdenii* and T2D, along with its association with medication treatment, consistent with prior research ^70,84,85^. However, the exact mechanisms underlying how *M. elsdenii* influences the development of T2D remain unclear and necessitate further investigation. Based on our data, we hypothesize that *M. elsdenii* may contribute to T2D development through the production of butyric acid. To validate this hypothesis, we conducted experiments on germ-free mice fed a high-fat diet. Our results confirmed that administering *M. elsdenii* prevented abnormal elevation of blood glucose levels in these mice. Furthermore, we observed an increase in butyric acid content in the intestines of mice treated with *M. elsdenii*. In conclusion, our results suggest that medication may elevate the levels of *M. elsdenii*, which in turn helps mitigate intestinal inflammation by producing short-chain fatty acids, thereby potentially addressing elevated blood glucose levels in individuals with T2D.

Overall, metaproteomics serves as a valuable tool for uncovering the roles of microorganisms in human health and their patterns of functional expression. Our data serve as a significant resource for investigating microbial functions, thereby enhancing our understanding of how microbes influence human health.

### Conclusion

We generated high-quality metaproteomic data from 1,967 fecal samples collected from a Chinese population. Our analysis delved into inter-individual variations of human proteins, microbial taxa, and gut microbiota functions. The findings underscore the effectiveness of population-based metaproteomics in representing actively functional biodiversity, elucidating significant variations in functional clusters, and offering a more reliable indicator of the temporal functional stability of the microbiome. We summarized highly expressed functions of each core microbe and pinpointed core microbes capable of expressing elevated levels of SCFA and vitamin B production. In addition, we investigated the correlation between 49 phenotypes and metaproteomic data. We outlined the microbial functional characteristics linked to T2D and highlighted how *M. elsdenii* alleviates T2D by generating butyric acid through lactate utilization. Our study offers valuable resources for investigating the functions of the gut microbiota.

## Acknowledgments

This study and the coauthors of this study are jointly supported by grants from the National Key R&D Program of China (No. 2021YFA1301600 and 2020YFE0202200), the National Natural Science Foundation of China (82073529, 81903316), and the Westlake Educational Foundation. We thank Westlake University Supercomputer Center for assistance in data analysis and storage, Dr Shu Zhu and his team for their advices.

## Author contributions

T. G., J.Z., and Y.C. designed and supervised the project. B.L. Z.M., E.C., M.Y., M.S., W.G., and K.Z. collected the samples and metadata. Shuang L., Y.S., E.C., M.Y., Y.X., Sainan L., Y.W., H.G., K.Z., J.Y., X.C., and X.W. generated the data. Shuang L., Y.S., Z.M., Z.X., P.L., M.S., W.G., and W.J. analyzed the data. Shuang L., Y.S., and T.G. drafted the manuscript with inputs from all co-authors. Shuang L., Y.S., Z.M., and B.L. contribute equally to this work.

## Competing interests

T.G. is a shareholder of Westlake Omics Inc. P.L., and J.Y. are currently employees of Westlake Omics Inc. The remaining authors declare no competing interests.

## Materials and Methods

### The GNHS cohort

Our study population included 1,967 stool samples from 1,399 participants from the GNHS cohort ^86^ of middle-aged and elderly people (40–75 years old) from the Guangzhou city urban area. Stool samples collected on-site at the Sun Yat-sen University were manually stirred, stored on ice, and transferred to −80 °C within 4 hours. Additionally, 25 physiologically measured or questionnaire-based phenotypic factors^87^ were collected along with samples. Specifically, metadata variables excepted antibiotic usage were classified into five phenotypic categories, containing demographics, lifestyle, anthropometrics, biochemical measures, and metabolic disorders. Also, the metadata types can be continuous (*e.g.*, age, body mass index, and blood lipid levels), binary (*e.g.,* sex, disease status), or categorical (*e.g.*, Bristol scale scores). More details, including the definition of metabolic syndrome, type 2 diabetes, and hypertension, as well as the statistics of each phenotype, are provided in Table S1.

### The FH cohort

Additional 104 individuals (52–83 years old) were recruited in Guangdong Province, China^54^. From these participants, we collected stool samples and their corresponding metadata as the GNHS cohort does. Detailed metadata information is provided in Table S1.

### Batch design

To minimize the batch effects, the 1,967 samples were randomly assigned to 96 batches by individuals’ age, gender, and sampling stage (baseline or follow-up). We designed intra-batch and inter-batch biological and technical replicates to monitor reproducibility during sample preparation and mass spectrometry (MS) acquisition. In each batch, two fecal samples were weighed twice and processed as intra-batch biological replicates. The inter-batch biological replicate samples mixed from randomly selected fecal samples were also designed in each batch before sample preparation. Also, after sample processing, peptides of two randomly selected fecal samples were injected twice for MS analysis in one batch as intra-batch technical replicates. The inter-batch technical replicates mixed from randomly selected peptides were also included in each batch. A batch contained 21 samples, two intra-batch biological replicates, two intra-batch technical replicates, one inter-batch biological replicate, and one inter-batch technical replicate. Altogether, we finally allocated 1,967 samples, 98 inter-batch technical replicates, 95 inter-batch biological replicates, 169 intra-batch biological replicates, and 185 intra-batch technical replicates to 96 batches (2,514 MS runs in total).

### Sample processing

For each sample, about 200 mg stools were gently resuspended in 500 μL cold phosphate buffer (PBS) and then centrifuged at 500 × *g* and 4°C for 5 min to collect the supernatant. This process was repeated twice to fully collect gut microbes and discard epithelial cells and food residues. Supernatants were combined (about 1.5 mL) and centrifuged at 500 × *g* and 4 °C for 10 min to discard more debris. Finally, supernatants (700 μL) were centrifuged at 18,000 × *g* and 4°C for 20 min to collect microbe pellets. Proteins were then extracted using 250 μL lysis buffer (4% w/v SDS and cOmplete Tablets (Roche) in 50 mM Tris-HCl, pH = 8.0) from the pellets. Microbe pellets added with lysis buffer were boiled at 95 °C for 10 min. After cooling at room temperature, samples were ultrasonicated at 40 Khz (SCIENTZ) for 1 h on ice. Following extraction, the lysis buffer containing sample proteins was centrifuged at 18,000 × *g* for 5 min for debris discarding. The resulting 200 μL supernatants were transferred into a new tube and added with a 5-fold volume of pre-cooled acetone. After vigorous shaking, proteins were precipitated overnight at −20 °C. Proteins were then pelleted by centrifuging at 1,000 × *g* for 3 min and washed with acetone three times at room temperature. The purified protein pellets were then completely re-dissolved in a dissolving buffer (8 mM urea and 100 mM ammonium bicarbonate). BCA protein assay kits were used to detect protein concentrations^55^. To prepare for the protein digestion, protein (50 μg) from each sample was reduced using 10 mM tris (2-carboxyethyl) phosphine (TCEP, Adamas-beta) and then alkylated with 40 mM iodoacetamide (IAA, Sigma-Aldrich). The first round of digestion was performed by adding 0.5 μg trypsin (Hualishi Tech) to the protein mixture, which was incubated for 4 h at 32 °C. Then, another 0.5 μg trypsin was added to the protein mixture, which was then incubated for 16 h at 32 °C. The tryptic peptides were desalted using solid-phase extraction plates (ThermoFisher Scientific, SOLAµ™) and then freeze-dried for storage. Dried peptides were resuspended in 2% acetonitrile, with 98% water and 0.1% formic acid (FA), before MS acquisition.

### Verification experiment of human proteins

Nine inter-batch biological replicate samples were used for the verification experiment. Firstly, the samples were divided into three groups (group 1, group 2, and group 3), each containing three samples as technical replicates. All samples were gently resuspended in 500 μL cold PBS and then centrifuged at 500 × *g* and 4°C for 5 min three times (group 1), four times (group 2), or five times (group 3) to collect the supernatant. The combined supernatants (about 1.5 mL, 2.0 mL, and 2.5 mL for group 1, group 2, and group 3) were centrifuged at 500 × *g* and 4 °C for 10 min to discard more debris (Fig. S3A). Next, 700 μL supernatants were centrifuged at 18,000 × *g* and 4 °C for 20 min to collect microbe pellets (Fig. S3A). The supernatant after centrifuging was collected into new tubes to detect the proteins that remained in the supernatant. Then, stool residues after microbe enrichment in group 1, group 2, and group 3 were further washed with 500 μL cold PBS and centrifuged at 500 × *g* and 4 °C for 5 min to collect the supernatant after microbe enrichment. The following procedures of sample preparation of microbe pellets were followed as what has been described above. The supernatants were directly added with a 5-fold volume of pre-cooled acetone to precipitate proteins. Then the proteins were digested and desalted as described above.

### High-pH reversed-phase fractionation

To generate a comprehensive spectral library of stool gut microbial proteins, randomly mixed peptides from 672 samples in GNHS cohort together with peptides from 11 healthy Chinese adults’ stool (five males and six females, their ages ranging from 22 to 33 years) were mixed into a library building pool (Fig. S1A). The pooled sample was then fractionated using high-pH reversed-phase liquid chromatography (LC). About 300 μg of purified peptides from the pool were separated using a nanoflow DIONEX Ultimate 3000 RSLC nano System (ThermoFisher Scientific) with an XBridge Peptide BEH C18 column (300 Å, 5 μm× 4.6 mm × 250 mm) (Waters) at 45 °C. The mobile phase of buffer A was water with 0.6% ammonia and pH = 10, while buffer B contained 98% acetonitrile and 0.6% ammonia, again with pH = 10. A 120 min gradient from 5% to 35% buffer B with a flow rate of 1 mL/min was applied. As a result, 120 fractions were collected at 1 min intervals and then combined into 30 fractions. Next, the fractions were freeze-dried and re-constituted in 2% acetonitrile, with 98% water and 0.1% FA.

### ddaPASEF mass spectrometry acquisition for library generation

The fractionated peptides were spiked with iRT (Biognosys)^88^ and loaded on a precolumn (5 μm, 100 Å, 5 mm × 300 μm I.D.) and then separated by a nanoElute UHPLC System (Bruker Daltonics) equipped with a 150 mm × 75 μm I.D. fused silica column custom-packed with 1.9 μm 120 Å C18 aqua (homemade) using a 95 min gradient at 300 nL/min. Buffer A contained 0.1% FA in water, while buffer B contained 0.1% FA in acetonitrile. The gradient flow was first set to 2% buffer B, linearly increased to 22% buffer B during the next 80 minutes, then linearly increased to 35% buffer B for additional 10 minutes, and finally increased to 80% buffer B during the next 2 minutes and kept at this level for another 3 minutes. A time-of-flight instrument coupled to a dual Trapped Ion Mobility Spectrometry (TIMS) analyzer (Bruker timsTOF Pro) equipped with a CaptiveSpray nano-electrospray ion source was used for MS acquisition. The TIMS section was operated with a 100 ms ramp time and a scan range of 0.6–1.6 V·s/cm^2^. One cycle was composed of 1 MS scan followed by 10 PASEF tandem-MS (MS/MS) scans. The MS and MS/MS spectra were recorded for mass-to-charge ratios (m/z) ranging between 100 and 1700. A polygon filter was used to filter out singly charged ions. For all experiments, the quadrupole isolation width was set to 2 Th for m/z < 700 and 3 Th for m/z > 800; between these values, the isolation width was linearly interpolated. The collision energy was ramped stepwise as a function of the increasing ion mobility: 20 eV energy start for the 1/K0 start (0.6 V·s/cm^2^) and 59 eV for the 1/K0 end (1.60 V·s/cm^2^). Eighty-seven MS profiles were acquired from fractionated peptides with repeated acquisitions. We also acquired one MS profile for the peptides of the library building pool. As a result, 88 LC-MS profiles were obtained for the spectral library building.

### diaPASEF mass spectrometry acquisition

A tandem system comprising a nanoElute UHPLC System (Bruker Daltonics) and a hybrid TIMS quadrupole time-of-flight mass spectrometer (Bruker timsTOF Pro) was used for MS measurements. For the LC analysis, 300 ng peptides were trapped at 500 Bar on the precolumn (5 μm, 100 Å, 5 mm × 300 μm I.D.) in 0.1 % FA/water and then separated along a 65 min LC gradient at a flow rate of 300 nL/min with a 150 mm × 75 μm ID fused silica column packed with 1.9 μm 120 Å C18. A linear gradient from 5 to 27% buffer B was applied for 50 min, followed by a second one from 27 to 40% buffer B for the next 10 min, and closed by a 2 min step to 80% buffer B, which was used for washing during the last 3 min. Buffer A: 0.1% FA in water; buffer B: 0.1% FA in ACN. The ion mobility range was limited to 0.7–1.3 V·s/cm^2^, and we applied four windows to each 100 ms diaPASEF scan. Fourteen of these scans covered the doubly and triply charged peptides’ diagonal scan line in the m/z ion mobility plane with 28 m/z narrow precursor windows. Other parameters were the same as those set during the ddaPASEF analysis.

### Spectral library construction

We used the integrated gut microbiome protein database described in a previous study^18^ and named it “IGC+”, to quantify microbial proteins. The IGC+ database contains the IGC 9.9 database^74^, fungi, virus sequences, and an in-house gut microbial gene catalog. The FragPipe platform (version 15), equipped with MSFragger (version 3.3)^89^, Philosopher (version 4.0.0)^90^, and EasyPQP (version 0.1.20), were chosen to build the spectral library. A three-step iterative search strategy for metaproteomics ^13,27^ was used with a few modifications. As detailed described in Fig. S1B, in the first step, we divided the IGC+ database into ten small databases and searched the 88 ddaPASEF files with a target-only version for database search. The matched protein sequences of the ten target-only searches for each ddaPASEF file were combined to reconstruct a simplified database for each MS profile (S1.fas–S88.fas). In the second step, each MS profile was searched against its corresponding simplified database using a target-decoy database search for accurate identifications. Proteins obtained from the 88 searches were aggregated into the concentrated database catalog of the GNHS fecal samples (GNHS.fas). In the third step, the 88 profiles were searched together against the GNHS database combined with a *Homo sapiens* protein catalog (Swiss-Prot, date 20211213) and the 245-contaminant database (MaxQuant, date 20160325) for the peptide to spectrum matches (PSMs) and peptide identifications. For the MSFragger analysis, the precursor and fragment tolerance were set to 20 ppm. Fragment ion series was set to b and y ions. Enzyme specificity was set to ‘trypsin’, allowing cleavages before proline. Up to two missed cleavages were allowed. The isotope error was set to 0/1/2. The peptide length was set from 7 to 50, and the peptide mass was set from 500 to 5000 Da. The carbamidomethylation of cysteine (57.021464 Da) was set as a fixed modification. The oxidation of methionine (15.99490 Da) and acetyl (Protein N-term) (42.010600 Da) was set as variable modification. The false discovery rate (FDR) was set to 0.01 for all the PSMs, ions, peptides, and proteins in the target-decoy searches. The final output from the iterative database search was imported into EasyPQP to establish the primary spectral library. The retention times of the peptides of each fraction were aligned using the iRT calibration. After obtaining the primary spectral library, we manually assigned peptides to proteins based on the maximum parsimony principle of protein inference (Fig. S2A). The final GNHS fecal spectral library included 506,470 peptides,1,433 unique human proteins with human-specific peptides, 121,702 unique microbial proteins with microbiome-specific peptides, and 18,143 microbial protein groups that comprising undistinguished microbial proteins.

### diaPASEF MS data analysis

The 2,514 diaPASEF profiles were searched against the final GNHS fecal spectral library using DIA-NN (version 1.8)^91^. The maximum mass accuracy tolerance was set to 10 ppm for both MS1 and MS2 spectra. The quantification mode was set to “Robust LC (high precision)”. The protein inference in DIA-NN was disabled. All the other settings were left to their default value. The output matrices, filtered at precursor q value ≤ 1% and global protein q value ≤ 1%, were used for metaproteomic analysis.

### Microbial functional annotation

The COGs were mapped by the microbial protein sequences with eggNOG-mapper (version 2.1.5) ^92^. The narrowest COG, KOG (Eukaryotic clusters of orthologous genes), and arCOG (Archaeal clusters of orthologous genes) of each protein were extracted. Proteins were also aligned to the KO annotation using GHOSTX from GhostKOALA (version 2.2) ^93^ with a bit-score threshold of 60. The best hit for each query was selected. Regarding microbial protein groups, we kept the microbial protein groups whose microbial proteins had the same functional annotations. When multiple functions were assigned to the same protein, all the functions were considered.

### Microbial taxonomic annotation

Taxonomic annotations were performed using the peptide-centric aligning software Unipept (version 2.2.1, searching date 2021.05.09) ^94^. The identified microbial peptides were cut according to the “Advanced missed cleavage handing” rules, and the cut peptides were further filtered to contain more than five and less than 50 amino acids. The filtered peptides were annotated using Unipept with the “Equal I and L” rule. The taxonomic assignment results were then manually filtered to the taxa of bacteria, archaea, fungi, or viruses. All the taxonomic annotations were filtered by at least two distinctive peptides, and their abundances were computed by summing the intensities of all the corresponding peptides. Within the taxonomic tree, the “noname” was added to the name of the narrowest taxon if there were no annotated taxa names in the middle taxon rank.

### Taxonomic and functional profiling of metagenomic samples

We also performed metagenomic shotgun sequencing for GNHS participants using the Illumina HiSeq platform (Illumina Inc.). We analyzed metagenomic sequences using a pipeline built as previously described^42^. We trimmed the sequences for quality using PRINSEQ (version 0.20.4) ^95^, and removed the human genome (*H. sapiens*, UCSC hg19) using Bowtie2 tool (version 2.2.5) ^96^. Taxonomic profiling was performed using MetaPhlAn2 (version 2.6.0) ^97^, which classifies metagenomic reads to taxa and yields their relative abundances in each sample. Functional profiling was performed by applying HUMAnN2 (version 2.8.1) ^98^. This analysis mapped reads to UniProt Reference Clusters (UniRef90) ^99^ and further grouped them into 9,381 COG groups.

### Analysis of richness, intensity, and BC dissimilarity

We mainly used three types of matrices for analysis, the human protein matrix, the microbial taxonomic composition matrices, and the microbial functional matrices. Three parameters were used to assess the differences between study individuals, levels of richness, intensity, and BC dissimilarity between samples. The BC distance was calculated using the function *vegdist* of the package vegan (version 2.5-7) in R.

### Quality control analysis

The data reproducibility was evaluated by the Spearman pairwise correlation and the BC distance between replicates. PCAs, performed with the command *prcomp* in R, was used to evaluate the batch effects of this study.

### Associations between matrices

The associations between matrices were done using the Mantel test. Specifically, the BC distance dissimilarity of the two matrices was inputted to calculate their Spearman correlation by the function *mantel* in vegan (version 2.5-7).

### Features’ accumulation curves and estimation of feature richness

The accumulation curves were plotted based on the observed richness of features with increasing numbers of collected samples. The observed richness was calculated in subsets of increasing size composed of randomly chosen samples (500 repetitions for each sample size). The estimated richness was calculated based on the Chao1 index using the function *poolaccum* from the package vegan (version 2.5-7) in R. When the samples reached a saturation number (*i.e.*, the observed richness was equal to the estimated saturation number and remained stable even with further increased sample number), we can assume these samples to cover the full diversity of the studied population.

### GO enrichment analysis

The human protein GO biological process enrichment was performed using the package clusterProfiler (version 4.2.0) in R. The package org.HS.eg.db (version 3.14.0) was used to annotate the human proteins. For statistically significant results, both the BH-adjusted *p-*value and the q-value were required to be lower than 0.05.

### Microbial function pathway enrichment

The pathway enrichment was performed using the R package MicrobiomeProfiler. For population-level pathway enrichment, the core KOs of metaproteomics and metagenomics were selected and enriched by enrichKO function with parameters minGSSize =10 and maxGSSize = 200. Statistically significant enriched pathways were screened with BH-adjusted p value ≤ 0.05, and fold enrichment value (GeneRatio / BgRatio) ≥ 2.

### Fecal butyric acid content analysis

The fecal butyric acid content was detected by targeted metabolomics through ultra-high-performance liquid chromatography-tandem mass spectrometry (UPLC-MS/MS) system (ACQUITY UPLC-Xevo TQ-S, Waters Corp., MA, USA). Detailed information about the measurements could be refered to the previously study (Jiang et al., 2020).

### Covariate analysis

The associations between each phenotypic factor and variation of human protein, gut microbiome function, or gut microbial taxon were determined by PERMANOVA on the human protein, metaprotein, KO, COG, species or genus levels (Bray-Curtis dissimilarities using the function *adonis* and the vegan R package, version 2.5-7). The *p-*value of each phenotypic factor was determined using 1,000 permutations and adjusted with the BH method. The phenotypic factors with adjusted *p* ≤ 0.05 were selected as covariates.

### Association analysis between individual features and the metadata

The generalized linear model (GLM) (implemented with the R package glm, version 4.1.1) was used to identify the association between the abundance of features (metaproteins, human proteins, COGs, KOs, species, and genera) and each phenotype. Only the features with higher than 10% prevalence in the GNHS cohort were used for the GLM analysis. Then the feature abundance data were firstly INT transformed to bring them all on the same scale. To avoid the covariate effects potentially associated with feature abundances, for the phenotypes excluded “Sex”, “Age” and “Bristol Scale”, we included these three metadata variables as the confounding covariates in the GLM models. In the GLM model for the three phenotypes, we use the other two metadata variables as the confounding covariates. For each *p-*value, multiple testing corrections were performed using the BH adjustment. The associations with q value (adjusted *p*) ≤ 0.05 were considered statistically significant.

### Machine learning

A total of 785 metaproteins, 23 human proteins, 268 KOs, 279 COGs, 12 species, and 14 genera that were related to T2D (GLM, q ≤ 0.05) were used for the metaproteomic-based T2D classification. The GNHS samples were randomly divided into the training set and internal validation set with a 4:1 ratio. Then we started to train the model using the training set. Firstly, we used the eXtreme Gradient Boosting (XGBoost) wrapped recursive feature elimination algorithm to select the features for our model at metaprotein, human protein, KO, COG, species, or genus level, respectively. We then counted the occurrence of each feature in the 100 candidate feature sets. We extracted the features with more than 70 occurrences as the final input feature set and feed it into an XGB model. We then fine-tuned the XGBoost model using the Genetic Algorithm hyperparameter searching with an early stop of three generations without AUC improvements. Finally, we tested the trained model in the internal validation set to check the generalization of the model. And an independent cohort (FH cohort) with 104 individuals was introduced for an independent test of the model.

### Co-occurrence Network

The linkages between the species and their corresponding functions were generated by their common peptide sequences, while the co-expressed linkages between intra-levels and inter-levels were calculated by the Spearman correlation (BH adjusted p-value ≤ 0.05, | r | ≥ 0.3 for inter-levels, | r | ≥ 0.8 for intra-levels).

### Data visualization

The R packages ggplot2 (version 4.1.3), ggpubr (version 0.4.0), gghalves (version 0.1.1), Goplot (version 1.0.2), forplo (version 0.1.0) and venn (version 1.10) were used for data visualization. The taxonomic tree was visualized using graphlan (version 1.1.3). The co-occurrence networks were visualized with the interactive platform Gephi (version 0.9.2). Additionally, we used the Fruchterman Reingold layout algorithm to visualize the networks. The modularity was built using the fast unfolding of communities in large networks. Different node colors represented different module communities. The size of the nodes was determined using the number of links between the nodes with other nodes.

### In vitro bacterial growth experiments

Precultures of *M. elsdenii* DSM 20460 were inoculated in anaerobic workstation (5% hydrogen, 10% carbon dioxide, and 85% nitrogen) as single colonies in modified reinforced clostridial medium (mRCM) medium. The medium contained the following components per liter: peptone (10g), beef extract powder (10g), yeast extract (3g), glucose (5g), fructose (10g), sodium chloride (5g), L-cysteine hydrochloride (0.5g), sodium acetate (3g), and resazurin (0.001g). After 24 hours of incubation, the preculture was inoculated into fresh mRCM broth at a concentration of 4% (v/v) and incubated for an additional 15 hours. The broth was supplemented with either 20 mM metformin, 20 mM acarbose, or 50 μm glimpiride in a 96-well microplate with 12 replicates per group. The impact of the drugs on bacterial growth was assessed by measuring the OD600 value of the bacterial culture and counting the number of live bacterial cells using BactoBox V7.2 (SBT Instruments), to ensure consistent results.

### Animal trail

Mouse experiments and all associated manipulations were carried out in accordance with the approved protocol by the research ethics committee of GNOTOBIO (permit JTAW20230705), following standard care and husbandry guidelines. Germ-free male C57BL/6 mice were randomly assigned to four ABSL-1 animal facilities with controlled environments. These environments maintained a 12-hour dark-light cycle (7 p.m. to 7 a.m.), 50±10% humidity, a temperature of 22 ± 2 °C, and provided ad libitum access to food and water. Following a 3-day acclimation period, the mice were divided into four experimental groups (n = 7). (1) Control group (CD group): These mice were fed a chow diet (catalog no. D12450J, Research Diets) with 10% of calories from fat, 20% from protein, and 70% from carbohydrates, providing 382 kcal per 100 g. They were administered 0.9% saline solution by gavage six times in the first two weeks. (2) High-fat diet group (HFD group): These mice were fed a high-fat diet (catalog no. D12492i, Research Diets) with 60% of calories from fat, 20% from protein, and 20% from carbohydrates, providing 524 kcal per 100 g. They were administered 0.9% saline solution by gavage six times in the first two weeks. (3) HFD+*M. elsdenii* group: These mice were fed a high-fat diet and administered *M. elsdenii* cells at a concentration of 10^9^ CFU by gavage six times in the first two weeks. (4) HFD+*L. reuteri* group: These mice were fed a high-fat diet and administered *L. reuteri* cells at a concentration of 10^9^ CFU by gavage six times in the first two weeks. After exsanguination, the mice were euthanized by cervical dislocation, and the intestines were dissected to collect the corresponding contents.

## Notes

### Summary of Updates

Author informations and affiliations updated;

